# Enemy or ally: a genomic approach to elucidate the lifestyle of *Phyllosticta citrichinaensis*

**DOI:** 10.1101/2021.11.27.470207

**Authors:** Valerie A. Buijs, Johannes Z. Groenewald, Sajeet Haridas, Kurt M. LaButti, Anna Lipzen, Francis M. Martin, Kerrie Barry, Igor V. Grigoriev, Pedro W. Crous, Michael F. Seidl

## Abstract

Members of the fungal genus *Phyllosticta* can colonize a variety of plant hosts, including several *Citrus* species such as *Citrus sinensis* (orange), *Citrus limon* (lemon), and *Citrus maxima* (pomelo). Some *Phyllosticta* species have the capacity to cause disease, such as Citrus Black Spot, while others have only been observed as endophytes. Thus far, genomic differences underlying lifestyle adaptations of *Phyllosticta* species have not yet been studied. Furthermore, the lifestyle of *Phyllosticta citrichinaensis* is ambiguous, as it has been described as a weak pathogen but Koch’s postulates may not have been established and the presence of this species was never reported to cause any crop or economic losses. Here, we examined the genomic differences between pathogenic and endophytic *Phyllosticta* spp. colonizing *Citrus* and specifically aimed to elucidate the lifestyle of *Phyllosticta citrichinaensis*. We found several genomic differences between species of different lifestyles, including groups of genes that were only present in pathogens or endophytes. We also observed that species, based on their carbohydrate active enzymes, group independent of their phylogenetic association, and this clustering correlated with trophy prediction. *Phyllosticta citrichinaensis* shows an intermediate lifestyle, sharing genomic and phenotypic attributes of both pathogens and endophytes. We thus present the first genomic comparison of multiple citrus-colonizing pathogens and endophytes of the genus *Phyllosticta*, and therefore provide the basis for further comparative studies into the lifestyle adaptations within this genus.

## 1 Introduction

Fungal and oomycete phytopathogens are a major threat to global food security (Fisher *et al*. 2012). Despite many technological and methodological developments, such as the development of disease-resistant crops, this threat remains a pressing concern for humankind due to emergence of new or adapted species, and a lack of in-depth understanding of disease mechanisms and their genomic basis (Fudal *et al*. 2009; Singh *et al*. 2011; Fisher *et al*. 2012).

Plant associated fungi and oomycetes can be broadly classified as pathogens, endophytes, or saprotrophs, i.e., they are classified based on their capacity to cause disease symptoms. Furthermore, these microbes can be linked to five different trophic classes based on their specific feeding behavior (Kabbage *et al*. 2015). Necrotrophs are characterized as pathogens that feed on dead tissue, biotrophs as pathogens that feed on living tissue, and hemibiotrophs are pathogens that go through an initial biotrophic phase before switching to a necrotrophic phase (Oliver and Ipcho 2004). In the same classification model, non-pathogenic species that live within a plant are classified as endophytes, while species that live only on decaying plant material are referred to as saprotrophs. This classification model, which is mainly based on observational data, clearly has limitations, for instance when one species is classified as a necrotroph when interacting with one host but as biotroph when interacting with another (Veloso and Van Kan 2018). Consequently, much research in recent years has focused on establishing the genomic basis underlying differences between species that exhibit different lifestyles. Uncovering these genomic signatures would provide a more reliable method of classification and an increased understanding of host colonization and disease mechanisms, which is of significant importance in developing more effective disease management strategies (Haridas et al., 2020; Möller and Stukenbrock, 2017; O’Connell et al., 2012; Ohm et al., 2012; Plissonneau et al., 2017; Spanu, 2012).

A common feature of various investigations into the genomic basis of pathogenicity is the identification of specific adaptations present in one lifestyle but absent or reduced in the other (Klosterman *et al*. 2011; Gardiner *et al*. 2012; Kim *et al*. 2016). A major current focus is the study of effectors, which are often defined as small secreted proteins that play an important role in establishing the interaction with the host, for instance by degrading the host cell wall or shielding the pathogen from detection by the host immune system (Rovenich *et al*. 2014; Lo Presti *et al*. 2015; Fouché *et al*. 2018). Effectors are often shared by strains and sometimes by species that colonize the same host (Chiapello *et al*. 2015; van Dam *et al*. 2017), and on rare occasions are even passed on to a separate species through horizontal gene transfer (Gardiner *et al*. 2012; van Dam and Rep 2017). The identification of known effectors and other genes that are present only in species of a specific lifestyle can therefore provide useful information when studying the genomic basis of pathogenicity (Gibriel *et al*. 2016). However, as hosts rapidly evolve mechanisms to recognize effectors to re-establish immunity, effectors frequently mutate resulting in rapid effector diversification to avoid detection by the host immune system. Thus, effector repertoires in separate fungal lineages may differ significantly (Rovenich *et al*. 2014; Lo Presti *et al*. 2015).

Carbohydrate active enzymes (CAZymes) play diverse roles in degradation and biosynthesis of carbohydrates such as those found in plant cell walls. For example, plant pathogens can utilize CAZymes to penetrate the host cell wall to establish symbiosis and to liberate carbohydrates from host tissues for growth and reproduction (van den Brink and de Vries 2011; Kubicek *et al*. 2019). Thus, CAZymes can also contribute to virulence, and differences in CAZyme repertoires can mediate microbial lifestyle differences (ten Have *et al*. 2002; King *et al*. 2011; Hane *et al*. 2020). Consequently, CAZymes have been used to propose new lifestyle classification models for oomycete and fungal species (Hane et al., 2020). For instance, Hane and colleagues recently proposed five new trophic classes based on primary nutrient source preferences as approximated by the presence and abundance of CAZymes directly predicted from genome assemblies (Hane et al., 2020): polymertrophs correspond best to necrotrophs, and have received their name due to a preference for polymeric carbohydrates as primary nutrient sources. In contrast, monomertrophs, which correspond best to symbionts and biotrophs, prefer monomeric primary nutrient sources. Mesotrophs are an intermediate group utilizing both monomeric and polymeric nutrient sources, and correspond best to hemibiotrophs. Vasculartrophs are similar to hemibiotrophs, but also include species commonly classified as wilts, anthracnoses, and rots. The saprotrophic class remains a separate group, encompassing species that feed mainly on decaying plant material. These new trophic classes were proposed based on a broad and phylogenetically diverse set of phytopathogenic fungi and oomycetes (Hane et al. 2020), including several members of *Dothideomycetes*, a diverse class including many plant-associated fungi as well as fungi adapted to other lifestyles such as marine or soil environments (Haridas *et al*. 2020).

Within the *Dothideomycetes*, members of the genus *Phyllosticta* are particularly well suited to study the genomic basis of lifestyle adaptation and phytopathogenicity. *Phyllosticta* contains at least 50 species that are able to associate with a broad range of plant hosts (Wikee *et al*. 2013b), but species that colonize *Citrus* are of particular interest as they comprise both endophytes and pathogens while being phylogenetically closely related (Wikee *et al*. 2013b; Guarnaccia *et al*. 2019). The most well-known species of this genus is *Phyllosticta citricarpa* which causes Citrus Black Spot, a disease causing significant economic losses worldwide and which therefore has a quarantine status in Europe (Kotzé 2000; European Food Safety Authority 2014; Eustáquio Lanza *et al*. 2018). *Phyllosticta paracitricarpa* is closely related to *P. citricarpa*, bears a strong morphological resemblance and appears to cause similar disease symptoms on citrus (Guarnaccia *et al*. 2017). Other pathogens include *P. citriasiana*, described from several citrus hosts in Asia, *P. citrimaxima*, described from *Citrus maxima* in Thailand, and *P. citrichinaensis*, described as a weak pathogen from several citrus hosts in China (Wulandari *et al*. 2009; Wang *et al*. 2012; Wikee *et al*. 2013b). Endophytic species within the genus are *P. capitalensis* with a very broad host range and present on all continents, *P. paracapitalensis*, known from citrus in Europe, and *P. citribraziliensis*, currently known only from citrus in Brazil (Glienke *et al*. 2011; Wikee *et al*. 2013a; Guarnaccia *et al*. 2017).

Genomes were recently published for *P. capitalensis*, *P. citriasiana*, *P. citribraziliensis*, *P. citricarpa*, *P. citrichinaensis*, and *P. paracitricarpa*, with genome sizes ranging between 29– 32 Mb (Guarnaccia *et al*. 2019). These genomes pave the way for comparative genomic studies aimed to disentangle lifestyle differences within the genus *Phyllosticta* (Guarnaccia *et al*. 2019). Although both mating types (MAT1-1 and MAT1-2) are reported to exist for both endophytic and pathogenic species (Guarnaccia *et al*. 2019; Petters-Vandresen *et al*. 2020), the pathogenic strains for which genomes have been published are all heterothallic and only of the MAT1-2 mating type. In contrast, the sequenced strain of the endophytic *P. citribraziliensis* is heterothallic and of the MAT1-1 mating type, while the other sequenced endophyte*, P. capitalensis*, is homothallic and therefore contains both mating types genes (Guarnaccia *et al*. 2017, 2019; Petters-Vandresen *et al*. 2020). *Phyllosticta citrichinaensis* is also homothallic, but the MAT1-2 idiomorph in *P. citrichinaensis* is present in a separate location from the mating type locus (Petters-Vandresen *et al*. 2020). This phenomenon sets *P. citrichinaensis* apart from the other species in the genus as the configuration of the mating type locus is typically very conserved amongst *Phyllosticta* species (Petters-Vandresen *et al*. 2020).

Previous comparative analyses between pathogenic and endophytic *Phyllosticta* spp. have been hampered by the quality of genomes and the availability of only a single endophyte genome (*P. capitalensis*), which was relatively distantly related to the species it was compared to, and consequently genomic adaptations towards these two broad lifestyles remained unclear (Wikee *et al*. 2013b; Rodrigues *et al*. 2019; Wang *et al*. 2020). Therefore, a comparison of new and high-quality genomes which includes multiple species of different lifestyles could provide the necessary foundation to finally discovering the genomic underpinning for phytopathology in *Phyllosticta*, which is essential for the development of better disease management strategies.

*Phyllosticta citrichinaensis* was originally described as a weakly aggressive pathogen of several citrus hosts in China as it was isolated from freckles or spots on fruits or leaves of citrus (Wang *et al*. 2012). However, lesions never exhibited typical pycnidia, the presence of this species was never reported to cause any crop or economic losses and Koch’s postulates may not have been established (Wang *et al*. 2012), and thus its lifestyle remains ambiguous. *P. citribraziliensis* is a very close relative of *P. citrichinaensis* and has been described only as an endophyte from Brazil. Therefore, if these two species were certain to be of two different lifestyles, these species would be ideal to study pathogenicity in *Phyllosticta*. As the genome of *P. citrichinaensis* has not been included in earlier comparative work focused on lifestyle differences (Rodrigues *et al*. 2019; Wang *et al*. 2020), a thorough study of its genome and comparison to the genomes of the other species in this genus could provide valuable information on this species’ lifestyle as well as genomic underpinning of disease mechanisms of other species in this genus. Here, we present the first comparative genomics study using multiple complete genomes of two endophytic and three phytopathogenic *Phyllosticta* species and established genomic differences between species of different lifestyles within this genus. In addition, we use these data in an attempt to elucidate the lifestyle of the ambiguous *P. citrichinaensis*.

## 2 Materials and Methods

### 2.1 Sequencing, annotation, genome quality and availability

All non-*Phyllosticta* genomes were previously published (Haridas *et al*. 2020) and are available on MycoCosm (https://mycocosm.jgi.doe.gov/Dothideomycetes; Grigoriev et al., 2014). The database identifiers (DBIDs) that are given by the Joint Genome Institute (JGI) to identify specific genomes, and which can be used to access the genome’s online portal (https://mycocosm.jgi.doe.gov/DBID, e.g., https://mycocosm.jgi.doe.gov/Aaoar1) are listed in Suppl. Table 1. Seven of the eight *Phyllosticta* genomes included in our analyses were also previously published (Guarnaccia *et al*. 2019), and are available on MycoCosm (mycocosm.jgi.doe.gov/Phyllosticta).

*Phyllosticta citrichinaensis* liquid cultures (250 ml Malt peptone broth in 500 ml Erlenmeyer flasks) were incubated at 25 °C and 180 rpm for 10 to 14 days, after which genomic DNA was isolated using the Qiagen Genomic-tip 100/G kit and the Qiagen Genomic DNA Buffer Set. The genome assembly of *P. citrichinaensis* genome (CBS 129764) was generated by the JGI using the PacBio long-read sequencing technology. Long-read sequencing data was assembled using Flye and the genome assembly was annotated using the JGI Annotation pipeline (Grigoriev *et al*. 2014). The genome assembly and annotation are available via the MycoCosm platform (https://mycocosm.jgi.doe.gov/Pcit129764). Quality assessments were performed using BUSCO 4.1.4 (Manni *et al*. 2021) and QUAST 5.0.2 (Gurevich *et al*. 2013) using default parameters. One-to-one whole-genome comparisons were performed using PROmer (default settings), which is part of the MUMMer 3.25 conda package (Marçais *et al*. 2018) and plotted with mummerplot using the --filter and --fat parameters. We used OrthoFinder 2.2.6 (Emms and Kelly 2019) to identify ortholog groups (OGs) across all 116 fungal genome annotations (Suppl. Table 2A). Ortholog groups unique to species of a specific lifestyles were identified using UpSetR (Suppl. Table 2B, Conway et al., 2017).

### 2.2 Secreted proteins and effectors

We used SignalP 5.0b Linux x86_64 (Almagro Armenteros et al., 2019) to predict secreted proteins in the predicted proteomes of all 116 fungal genomes, and subsequently applied EffectorP 2.0 (Sperschneider *et al*. 2016) to predict effectors within the secretomes. We visualized the distribution of OGs of which 50% or more of the genes were predicted to be a secreted protein or an effector, by generating a clustered heatmap in R using the ComplexHeatmap package (Gu *et al*. 2016).

### 2.3 Carbon utilization and CATAstrophy

Carbon growth studies were performed as described previously (Buijs *et al*. 2021). In short, 1-mm-diameter plugs from 2-week-old colony edges of *Phyllosticta* species were inoculated on 35 different carbon sources and incubated at 25 °C until the largest colony reached the edge of a 35-mm-diameter plate. As different *Phyllosticta* species demonstrate different growth speeds, this moment fell on different days after inoculation (between five and ten days). When the largest colony of a species reached the edge of a plate, colony diameters were measured on all sources and images were taken using a standard camera setup. This approach was chosen to be able to compare species with different growth speeds. All growth studies were performed in duplicate. Measurements were averaged and used to generate a clustered heatmap using the ComplexHeatmap package (Gu *et al*. 2016) in R (R Core Team, 2021).

We used CATAstrophy to predict lifestyles from CAZyme repertoires (Hane *et al*. 2020). To this end, we first used hmmpress to generate a local HMMER database of dbCAN 8 (Zhang *et al*. 2018). We then queried all 116 predicted proteomes with the local dbCAN HMMs database using hmmscan with the –domtblout parameter to create a domain table for each proteome. CATAstrophy was then ran on all 116 domain tables using parameters -p, -c, --model v8 and --format hmmer_domtab. The heatmap was created by identifying all OGs (as created previously using OrthoFinder) that contained a CAZyme, counting the number of genes in each OG for each species, and then generating a heatmap using the ComplexHeatmap package in R. Empty columns, e.g. CAZyme families for which no genes present in these species, were filtered out of the heatmap for visualization purposes, but are present in the original data (Suppl. Table 5).

## 3 Results

### 3.1 *Phyllosticta* genome assemblies are of high quality

Lifestyle differences are often driven by genomic adaptations (Ohm *et al*. 2012; Kabbage *et al*. 2015; Haridas *et al*. 2020), and we hypothesize that this also applies to *Phyllosticta* species with different lifestyles. Taxonomically, *Phyllosticta* belongs to *Dothideomycetes*, a fungal class with extensive genomic resources (Wikee *et al*. 2013b; Haridas *et al*. 2020). To enable studies in lifestyle differences in *Phyllosticta,* we made use of eight *Phyllosticta* genome sequences, seven of which were assembled and published previously (Guarnaccia *et al*. 2017, 2019). Here, we performed genome sequencing of *P. citrichinaensis* (CBS 129764), which is the second genome of this species to be sequenced, thereby enabling us to also evaluate intra-species variation. We included two genome assemblies of *P. citrichinaensis*, as well as the genome assemblies of the endophyte *P. citribraziliensis*, the closest relative of *P. citrichinaensis*, those of the two pathogenic species *P. citricarpa* and *P. paracitricarpa*, which are very closely related (Fig. 1A, also see Guarnaccia et al., 2019), of the pathogenic species *P. citriasiana*, and of the endophyte *P. capitalensis*, which is phylogenetically the least related to the other *Phyllosticta* species (Fig. 1A). As these genome assemblies include multiple species of both lifestyles (pathogens and endophytes), comparative genomics may reveal the genomic underpinning for lifestyle adaptations within this genus, and ultimately aid in determining the lifestyle of *P. citrichinaensis*.

**Figure 1.**
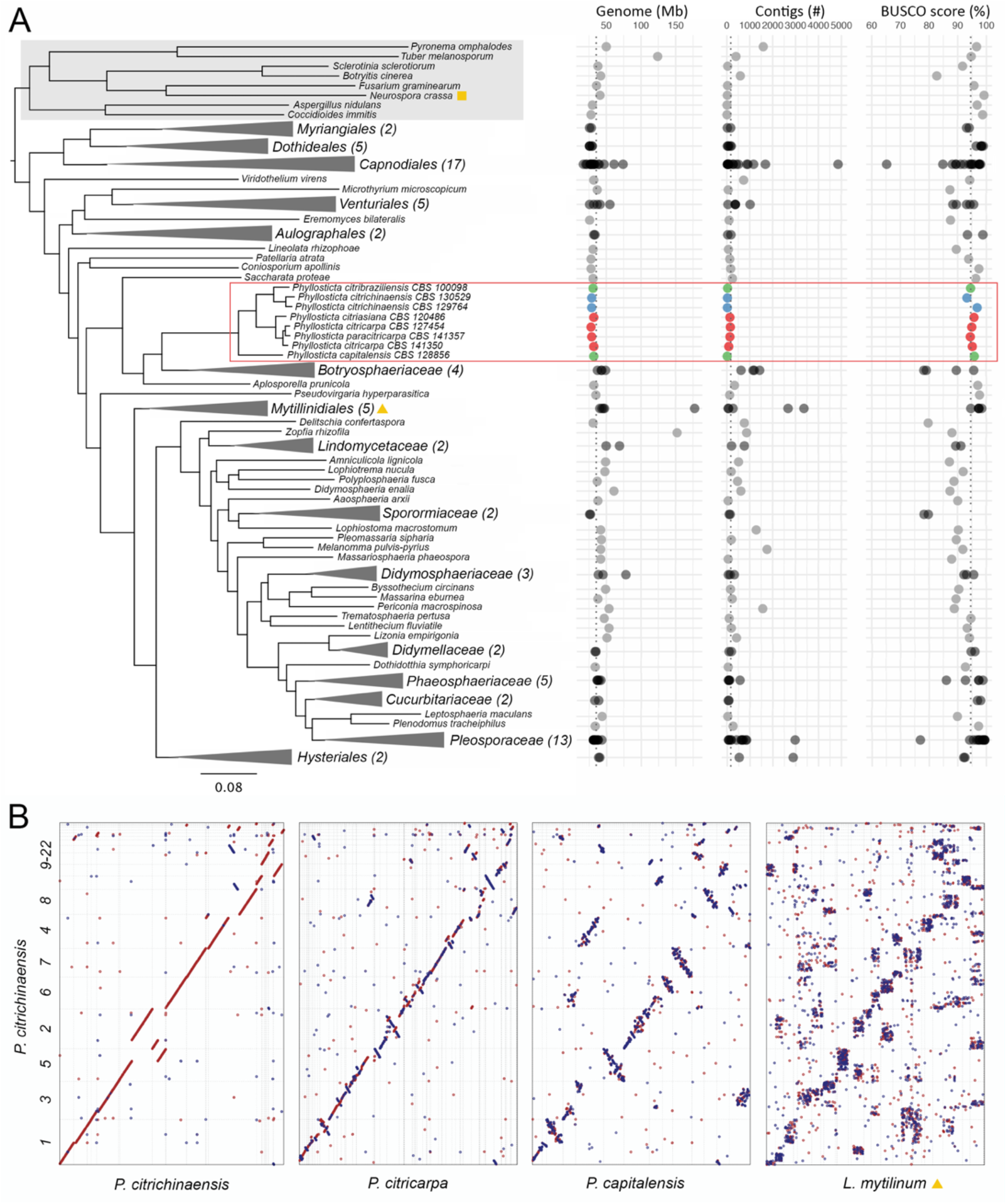
*Phyllosticta* genomes are of high quality and show mesosynteny. **A.** The phylogenetic relationship of more than 100 fungal species (116 genomes) is shown. The phylogenetic tree was generated using OrthoFinder, and sub-trees were collapsed manually to enhance readability. Grey box indicates species outside the *Dothideomycetes*, red rectangle indicates the genus *Phyllosticta*. Red orbs = *Phyllosticta* pathogens, green orbs = *Phyllosticta* endophytes, blue orbs = *P. citrichinaensis.* Yellow triangle and square indicate the locations of *Lophium mytilinum* and *Neurospora crassa*, respectively. **B.** Whole-genome alignments between *P. citrichinaensis* and *P. citricarpa, P. capitalensis*, and *L. mytilinum*, respectively, were generated using PROmer; the red line/dot = homologous area and the blue line/dot = reversed homologous area.

To determine genome assembly size, fragmentation, and annotation completeness of the *Phyllosticta* genomes, we used QUAST (Gurevich *et al*. 2013) and BUSCO (Manni *et al*. 2021), and compared the results to 100 previously published *Dothideomycete* genomes as well as eight genomes of fungal species outside of *Dothideomycetes* (Fig. 1A; Suppl. Table 1, Haridas et al., 2020). Compared to the other fungal species, *Phyllosticta* genome assemblies are of a slightly smaller size (29–32 Mb) as opposed to an average of 40 Mb in other *Dothideomycetes* (Fig. 1A). The *Phyllosticta* genome assemblies have a low number of contigs, namely 14 to 152 contigs compared with on average 471 contigs, and BUSCO scores between 93.3% and 95.8%, suggesting that the *Phyllosticta* genome assemblies are *en par* or above average quality in terms of genome contiguity and completeness compared to the other *Dothideomycetes* genomes, which should facilitate further comparative analyses.

### 3.2 *Phyllosticta* species show mesosynteny to species within the *Dothideomycetes*

*Dothideomycetes* are known to show different patterns of genome conservation as compared to other fungal classes: intra-chromosomal rearrangements lead to a seemingly random reshuffling of gene content within individual chromosomes, while genes themselves remain well conserved (Hane *et al*. 2011; Ohm *et al*. 2012), a pattern that has previously been named mesosynteny (Hane *et al*. 2011). To study the gene order conservation in *P. citrichinaensis* and in other *Phyllosticta* species compared with other fungal genomes, we performed whole-genome alignments of *P. citrichinaensis* to all other *Phyllosticta* and to two more distantly related fungi (Fig. 1B). When comparing the two *P. citrichinaensis* strains, we observed a clear pattern of macrosynteny with only a few inversions and translocations (Fig. 1B); a similar pattern could also be observed when comparing *P. citrichinaensis* to *P. citricarpa* (Fig. 1B) and *P. citribraziliensis* (Suppl. Fig. 1A). Interestingly, when comparing *P. citrichinaensis* to *P. capitalensis*, which is a more distantly related member of this genus (Fig. 1A), we observed an increased frequency of chromosomal rearrangements (mainly intra-chromosomal inversions with only few inter-chromosomal translocations), suggesting that these species are mesosyntenic (Fig. 1B). As expected, when moving (phylogenetically) further away from *P. citrichinaensis* and comparing its genome sequence to the one of *Lophium mytilinum*, a relatively obscure fungus from the order *Mytilinidiales* (*Dothideomycetes*) (Fig. 1A), a clear pattern of mesosynteny can be observed (Fig. 1B). When comparing *P. citrichinaensis* to *Neurospora crassa*, a species from the class *Sordariomycetes*, we only observed weak mesosynteny with the pattern apparently dissipating (Suppl. Fig. S1). Thus, our results clearly demonstrate gradually changing patterns of mesosynteny across progressively more distantly related species within the *Dothideomycetes*, but possibly also outside of this taxonomic class. Furthermore, we show that species within the genus *Phyllosticta* spur the mesosyntenic mode of evolution that has previously been described for other *Dothideomycetes* (Hane *et al*. 2011; Ohm *et al*. 2012).

### 3.3 *Phyllosticta* differ in gene number and functional annotation in a lifestyle dependent manner

Species of similar lifestyles often share (groups of) genes (Lo Presti *et al*. 2015; Kim *et al*. 2016). Consequently, we hypothesized that the presence or absence of specific genes may provide information about the (predominant) lifestyle of *P. citrichinaensis*. To be able to compare gene content over different species and strains, we used OrthoFinder (Emms and Kelly 2019) to identify ortholog groups (OGs) across all 116 predicted proteomes. Orthofinder identified 35,379 OGs containing 88.1% of all genes (Suppl. Table 2A). The 11.9% of genes that were not assigned to any OG likely constitute species-specific genes, which is to be expected given the taxonomically diverse set of fungal species considered in our study (Fig. 1A). In total, 1,794 OGs (5.1%) contained genes from all species, representing a fungal core genome. Of all OGs, 32.2% (11,352) contained at least one gene from a *Phyllosticta* species, and of those, 57.8% (6,558, Fig. 2) contained genes from all *Phyllosticta* species and 33.2% (3,764) were unique to *Phyllosticta* (i.e., they only contained *Phyllosticta* genes). The latter percentage is quite high because of the close taxonomic relation of some of the genomes: a rather large fraction of the *Phyllosticta* unique OGs contain two or three genes (2,734, 72.6%), as these often contain one gene from each of the two *P. citricarpa* genomes and one more gene from *P. paracitricarpa*. Genes that are unique to a species, i.e., sequences that are sufficiently different from other sequences, do not form a separate OG on their own and consequently are not considered in these statistics. Since the *P. citricarpa* and *P. paracitricarpa* genomes are so closely related, many of their “unique genes” are assigned to an OG, which causes the fraction of *Phyllosticta* unique OGs to be quite large.

**Figure 2.**
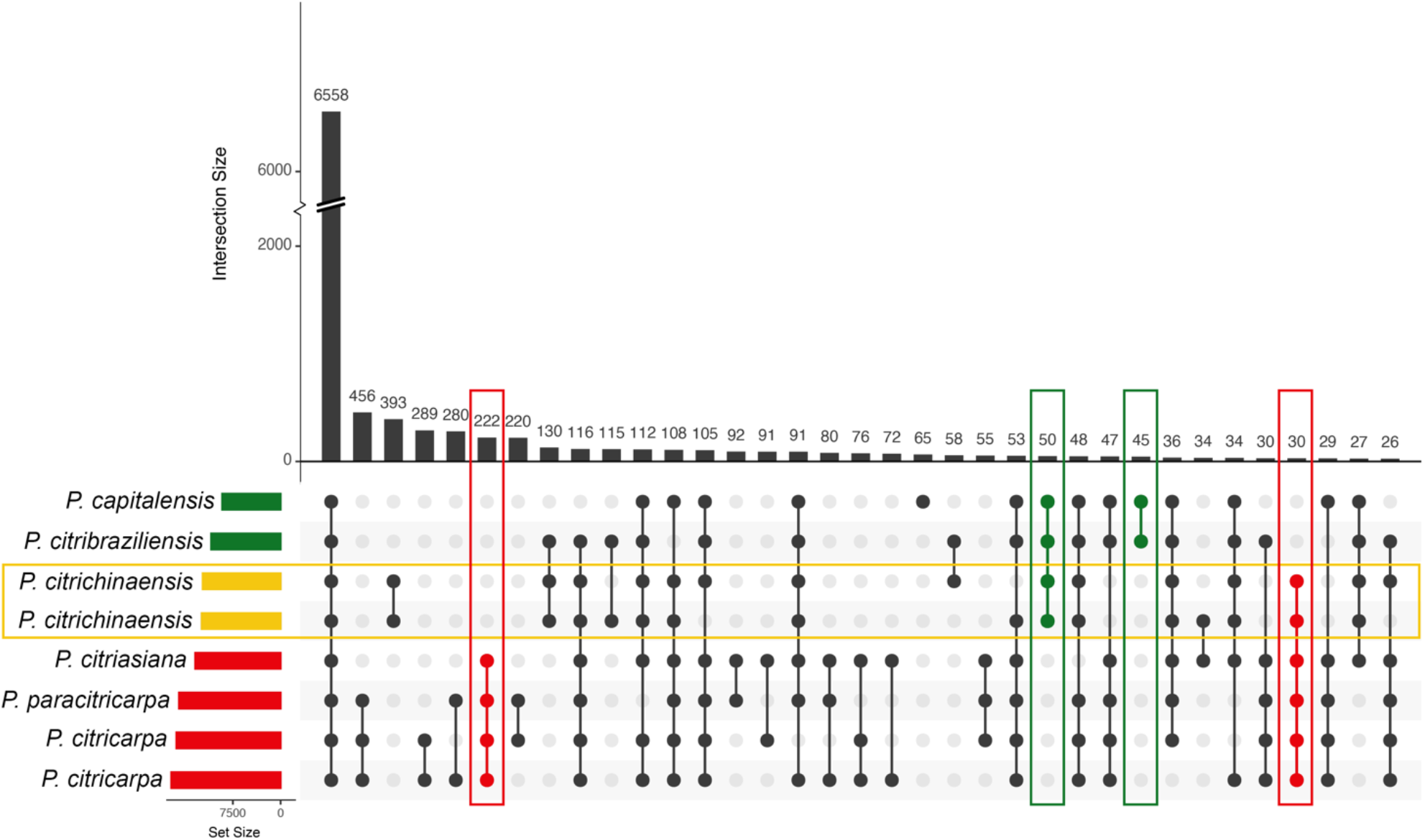
*Phyllosticta citrichinaensis* shares more unique OGs with endophytes than with pathogens, but pathogens share more OGs with each other. Yellow rectangles = *P. citrichinaensis*, red rectangles = pathogens, green rectangles = endophytes. Figure generated using UpSetR.

To discover lifestyle-associated genes in *Phyllosticta*, we compared OG content across the eight *Phyllosticta* species and sought to identify differences between species of different lifestyles. The number of OGs that is shared by *Phyllosticta* species of a specific lifestyle (45–222, Fig. 2 and Suppl. Table 2B) is much smaller than the total number of OGs that all species in this genus share (6,558). The OGs shared by *Phyllosticta* spp. of a specific lifestyle might also contain genes from species outside of this genus. *Phyllosticta* pathogens shared 222 OGs that are not present in *Phyllosticta* endophytes, a much higher number than the total of 45 OGs that are shared by endophytes and not present in pathogens, which is likely due to the larger phylogenetic distance between the two endophytes compared with the pathogenic *Phyllosticta* species (Fig. 1A). In addition to OGs that are unique to species of a specific lifestyle, OGs that are present in all species but differ in their abundance in species that share a lifestyle (e.g., there are more genes in species of one lifestyle compared to the others) may provide information on how species adapt to their lifestyles. We thus identified OGs for which the average number of genes in species of one lifestyle was higher than the average number of genes for species of the other lifestyle; we did not consider OGs that contained large outliers, i.e., one species having a much larger number of genes as compared to the other species. This resulted in a total of 87 OGs: 73 OGs that had more genes in endophytes and 14 OGs that had more genes in pathogens (Suppl. Table 2C), suggesting that achieving the endophytic lifestyle requires additional genes.

As functions of genes in OGs that are unique to, or enriched in, species of a certain lifestyle may underly lifestyles adaptations, and this may help to uncover which lifestyle *P. citrichinaensis* has, we looked into the annotations of the lifestyle-related OGs. In addition to the annotations of *Phyllosticta* genes, we also included the annotations of 50 out of the 108 species outside the genus *Phyllosticta* for which annotation data was available on the JGI database. The number of OGs in which at least one gene was functionally annotated (other than as hypothetical or expressed protein) varied widely between different lifestyle-related groups: while nearly 50% of the OGs that had more genes in endophytes received an annotation, less than 3% of the pathogen-only group did (Suppl. Table 3A–F). We further divided the individual functional annotations into KOG-classes (from the EuKaryotic Orthologous Groups tool, https://mycocosm.jgi.doe.gov/help/kogbrowser.jsf), which provide a high-level classification system to group genes with comparable activities. Of all KOG-classes, the class ‘Secondary metabolites biosynthesis, transport and catabolism’ was most often found to be associated with lifestyle-related OGs: only two of six lifestyle-related groups did not contain genes in this class, suggesting this group of genes may be useful to distinguish species of different lifestyles (Suppl. Table 3I).

To study secondary metabolite biosynthesis genes in more detail we used antiSMASH (Blin *et al*. 2019) to identify biosynthetic gene clusters (BGCs) in the eight *Phyllosticta* genomes. The total number of predicted BGCs varied from 20 in *P. paracitricarpa* to 24 in *P. citricarpa*, with no apparent differences between species of different lifestyles. Interestingly, one of the terpene clusters was predicted as a squalestatin in all pathogenic species, while it received no functional prediction in the endophytic species or in *P. citrichinaensis*, suggesting there is a difference in this cluster between species of different lifestyles. Squalestatins are predicted to be inhibitors of squalene synthase, which produces squalene, a sterol biosynthetic intermediate that is reported to play a role in mediating interactions between fungi and their plant hosts (Lindo *et al*. 2020). Therefore, further characterization of this BGC and others in *Phyllosticta* will be worthwhile for future studies into pathogenicity of *Phyllosticta* species.

Ortholog groups with more genes in *Phyllosticta* endophytes were more often functionally annotated, suggesting that these are generally better characterized and likely evolutionary conserved. We did not find any particular KOG-class to be annotated in higher abundance in endophytes, but nonetheless found a few interesting annotated OGs, such as six OGs that were annotated to belong to the ‘carbohydrate transport and metabolism’ class including a CAZyme family (GH55) gene and several transporters, suggesting a role for carbohydrate transport in lifestyle (Suppl. Table 3A–F, I). One endophyte-only OG contained the MAT1-1 gene, which was the result of all sequenced pathogenic strains having MAT1-2 mating types (Petters-Vandresen *et al*. 2020). In addition, although the *P. citrichinaensis* MAT1-1 gene is not present in this OG, we did find the MAT1-1 gene in the *P. citrichinaensis* genome assembly and found it to be highly similar to that of *P. citribraziliensis*, its closest relative.

The pathogen-only group was poorly functionally annotated; out of 222 OGs, only five received a functional annotation. The fact that such a large fraction of OGs could not be assigned a functional annotation is of interest as some of the non-annotated OGs in the pathogen-only group could contain putative effectors or genes that are otherwise involved in virulence, as effectors often remain unannotated in standard annotation pipelines (Sperschneider *et al*. 2015). In addition, two OGs received functional annotations that have previously been implied to be virulence factors and could therefore be interesting targets for future functional studies; a pectin lyase fold (Yang *et al*. 2018), and a cytochrome p450 (Siewers *et al*. 2005; Shin *et al*. 2017).

### 3.4 *Phyllosticta citrichinaensis* shares more lifestyle-specific OGs with endophytes, but follows an intermediate pattern in other lifestyle-associated OGs

The lifestyle of *P. citrichinaensis* is currently ambiguous (Wang *et al*. 2012), but the number of OGs that it shares with species of either lifestyle may provide clarity not only about the lifestyle of *P. citrichinaensis* itself but also about the differences between species of different lifestyles within this genus. The two *P. citrichinaensis* strains share more lifestyle specific OGs with endophytes (50) than they do with pathogens (30). In addition, the number of OGs that is shared by only the two endophytes and *P. citrichinaensis* is larger than the number of endophyte-only OGs that are not shared with *P. citrichinaensis* (Fig. 2, green rectangles), suggesting that *P. citrichinaensis* indeed compared well to endophytic species. However, for 30 out of 87 OGs that contained more genes in either pathogenic or endophytic species, the *P. citrichinaensis* gene numbers corresponded best to the endophytic numbers, while in 32 OGs, they corresponded to the pathogenic numbers. In 25 OGs, the number of genes of *P. citrichinaensis* corresponded to neither lifestyle (Suppl. Table 2C). As opposed to the numbers of OGs specific to species of one lifestyle, where *P. citrichinaensis* shared more with endophytes (Fig. 2), the numbers of *P. citrichinaensis* genes in OGs that contained more genes in either pathogenic or endophytic species thus indicate that it compares equally well to species of either lifestyle. Thus, this data suggests that presence/absence and/or gene abundance differences are not sufficient to provide insights into the lifestyle of *P. citrichinaensis*.

While we did not observe clear patterns in the types of functions that *P. citrichinaensis* shares with either endophytes or pathogens, we nevertheless could observe some interesting functional patterns in *P. citrichinaensis* (Suppl. Table 3G–H). For instance, two groups were annotated as heat shock proteins: an Hsp40 (DNAJC17) that had more genes in pathogens (3 in pathogens vs 1–2 in endophytes) as well as an Hsp70 that had more genes in endophytes (4–5 in pathogens vs 6–7 in endophytes). *Phyllosticta citrichinaensis* contains fewer genes in both groups, suggesting it may respond differently to stress. Furthermore, one group that had more genes in pathogens as well as in *P. citrichinaensis* (1 in endophytes versus 2–3 in pathogens) contains genes annotated as peroxiredoxin-1 or peroxiredoxin-6. Peroxiredoxins are necessary for full virulence in several fungal pathogens such as *Magnaporthe oryzae* and *Aspergillus fumigatus* as they offer an antioxidant defense against reactive oxygen species produced by the host as part of host defense responses (Mir *et al*. 2015; Rocha *et al*. 2018). It is thus possible that these additional genes in *Phyllosticta* pathogens and in *P. citrichinaensis* contribute to their virulence.

### 3.5 There is little difference between *Phyllosticta* endophytes and pathogens in the numbers of putative secreted proteins and putative effectors

Secreted proteins, including effectors, play an important role in lifestyle and virulence (de Wit *et al*. 2009; Lo Presti *et al*. 2015; Plissonneau *et al*. 2017). Based on the comparison of OGs in species of different lifestyles, we concluded that pathogenic species contain a large number of unannotated genes in pathogen-specific OGs (217 OGs). We hypothesized that some of these genes may be effectors or other secreted proteins, and that the presence of putative secreted proteins and effectors in the genomes of species may be an indicator for lifestyle differences. We therefore assessed the presence of OGs that contain 50% or more putative secreted or effector proteins in all 116 species used in this study. A total of 3,942 OGs (11% of 35,379 OGs) consisted of at least 50% secreted proteins (Fig. 3A, Suppl. Table 4A) and of these, 1,048 OGs consisted of at least 50% effectors (Fig. 3B, Suppl. Table 4B). Notably, only 62 OGs containing putatively secreted genes were present in all 116 species, and only one OG containing putative effectors contained genes from all 116 species, corroborating that secreted proteins and effectors are typically not shared between different species and especially effectors are rather species and/or strain specific (Stergiopoulos *et al*. 2012).

**Figure 3.**
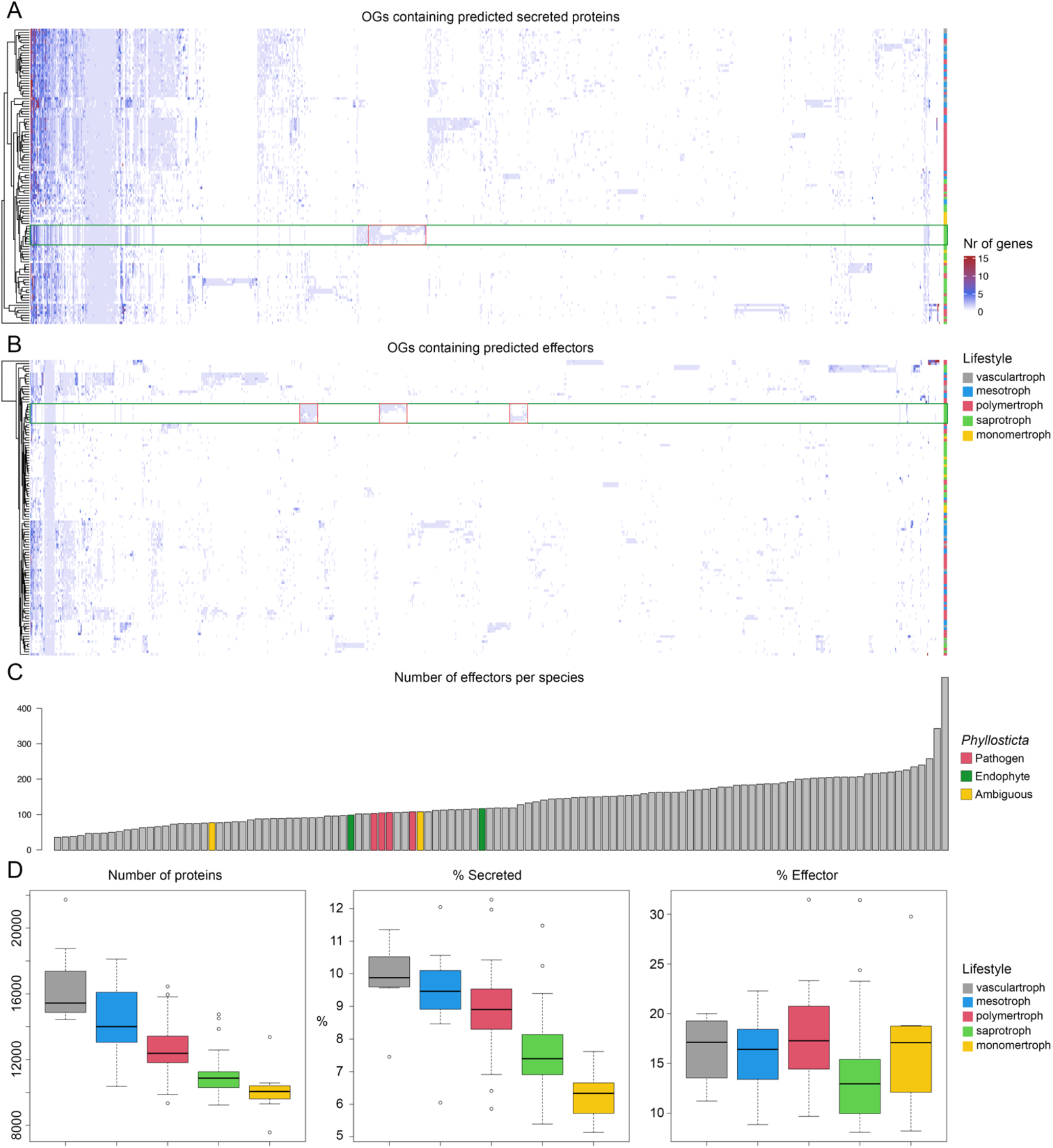
Species cluster independent of their taxonomic relationship based on presence of secreted or effector proteins. Clustered heatmaps of the number of genes in OGs that were predicted to contain **A.** secreted proteins and **B.** effectors. Lifestyles as predicted by CATAstrophy (Fig. 4). *Phyllosticta* species are highlighted with green rectangles, *Phyllosticta*-unique OGs are highlighted in red rectangles. **C.** Number of effectors per species (both those in OGs and singletons). *Phyllosticta* species are highlighted in red (pathogens), green (endophytes), and yellow (*P. citrichinaensis*). **D.** The total number of proteins, the percentage of proteins that is predicted to be secreted, and the percentage of secreted proteins that is predicted to be an effector, in species of different lifestyles (as predicted by CATAstrophy, Fig. 4).

*Phyllosticta* genes were present in a total of 762 putatively secreted OGs, with 432 of those containing genes from all *Phyllosticta* species included in this study. With an average of 786 genes per species, *Phyllosticta* species contain less genes in OGs encoding secreted proteins compared with other *Dothideomycetes* (an average of 1,033 genes, Suppl. Table 4). In addition, *Phyllosticta* species contained on average 110 putatively secreted genes that were not in an OG or that were the only protein predicted to be secreted in an OG (singletons), which is also less than we observed for the overall average of 128 singleton secreted genes (Suppl. Table 4). When assessing the total number of predicted secreted genes (both in OGs and singletons), *Phyllosticta* endophytes have more putative secreted proteins (average of 913) as compared to *Phyllosticta* pathogens (average of 897). However, when considering these as a percentage of the total predicted proteome size, this difference becomes negligible, with endophytes having a slightly smaller percentage of proteins that is secreted (7.89%) as compared to pathogens (8.01%, Suppl. Table 4C). We found 247 putatively secreted OGs to be unique to *Phyllosticta* (Fig. 3A), which mainly represent genes encoding proteins without predicted functions such as those often found to be putative effectors (Suppl. Table 4G, Lo Presti *et al*. 2015). Namely, only four out of 247 OGs contained genes that were functionally annotated, and interestingly, three out of four were predicted to function in carbon utilization (Suppl. Table 4G).

*Phyllosticta* species contain less genes in OGs encoding effectors with an average of 74 genes compared to the overall average of 93 genes. In addition, *Phyllosticta* species contained on average 30 putative effector genes that were not in an OG or that were the only effector in an OG (singletons), which is lower than the average of 44 singleton effector genes for the other *Dothideomycetes*. Comparable to secreted proteins, when assessing the total number of predicted effector genes (both in OGs and singletons), *Phyllosticta* endophytes appear to have slightly more putative effector genes (average of 109) as compared to *Phyllosticta* pathogens (average of 106.5), but when taken as percentage of the total number of predicted proteomes per species, this difference is negligible (0.95% vs 0.94%) (Fig 3B–D, Suppl. Table 4D). A total of 30 effector OGs contained genes from all *Phyllosticta* species while also containing genes from other species. One of these had higher gene numbers in endophytes, but received no annotation, while none had higher gene numbers in pathogens. Effector genes are often hypothesized to be species and/or strain specific (Lo Presti *et al*. 2015; Sperschneider *et al*. 2015). We identified in total 67 OGs containing effectors that are unique to *Phyllosticta*, seven of which were present in all pathogens but not in endophytes, two were present in all endophytes but not in pathogens, and one had higher gene numbers in endophytes (Suppl. Table 4E). Genes of the seven pathogen-only OGs were not colocalized as described in some fungal phytopathogens (Ma *et al*. 2010; Dong *et al*. 2015) but rather are spread across separate scaffolds. None of these unique effector genes were functionally annotated, suggesting that these have yet undescribed functions.

The occurrence of *P. citrichinaensis* effector genes in lifestyle-associated OGs could provide further evidence for its lifestyle. One of the seven effector OGs that only occurred in pathogens contained a gene from one of the *P. citrichinaensis* strains. In contrast, both endophyte-only effector OGs contained genes from *P. citrichinaensis*, in one case only from one strain, and in the other case from both strains. In the effector OG that had more genes in endophytes, *P. citrichinaensis* followed an intermediate pattern: one strain contained the same number of genes as pathogens, while the other contained the same number as endophytes. These data thus suggest that *P. citrichinaensis* follows an intermediate lifestyle.

### 3.6 *Phyllosticta citrichinaensis* clusters with pathogens based on carbon growth data

CAZymes enable fungi to utilize different carbon sources (van den Brink and de Vries 2011; Lombard *et al*. 2014), and are thought to be involved in fungal pathogenicity (ten Have *et al*. 2002; King *et al*. 2011; Kubicek *et al*. 2014; Hane *et al*. 2020). We have previously shown that carbon utilization capabilities differ between *Phyllosticta* spp. and uncovered a clear distinction in the ability of pathogens and endophytes to grow in the presence of sugar beet pulp; while the growth of endophytes was unchanged, pathogens were strongly inhibited (Buijs *et al*. 2021). To assess if *P. citrichinaensis* displays similar growth behavior to pathogens or endophytes, we grew *P. citrichinaensis* on 35 different carbon sources including sugar beet pulp (Fig. 4). Interestingly, growth of *P. citrichinaensis* is not inhibited by the presence of sugar beet pulp (Fig. 4A), suggesting that *P. citrichinaensis* behaves comparable to endophytic *Phyllosticta* spp. To further substantiate this observation, we performed hierarchical clustering of seven *Phyllosticta* strains based on their growth on all 35 carbon sources. Unanticipatedly, *P. citrichinaensis* clustered together with the pathogenic species rather than with the endophytes (Fig. 4B). Thus, although *P. citrichinaensis* is not inhibited by sugar beet pulp, it generally displays carbon utilization capabilities comparable with pathogens.

**Figure 4.**
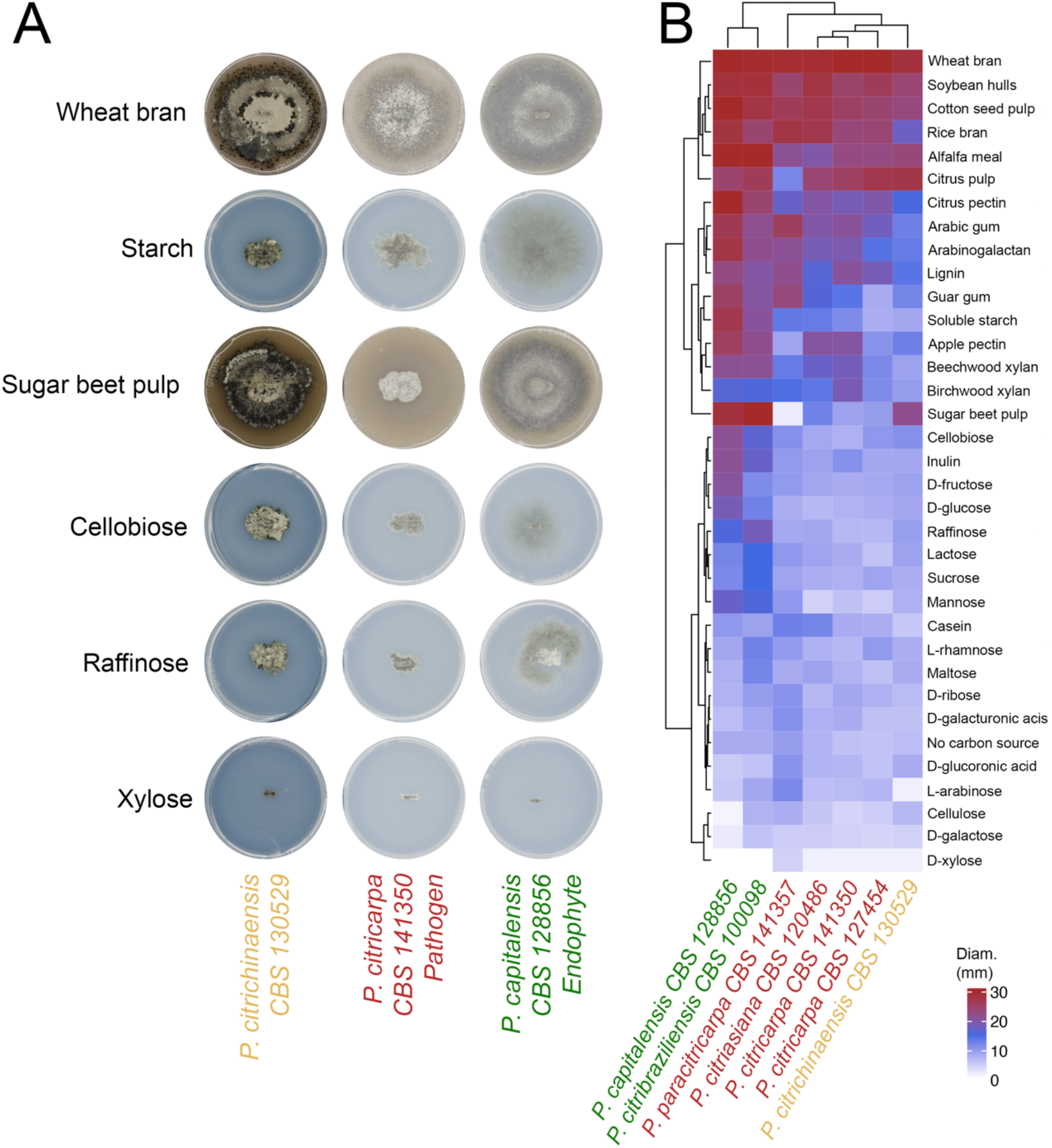
*Phyllosticta citrichinaensis* clusters with pathogens based on growth on 35 carbon sources, but behaves like an endophyte in the presence of sugar beet pulp. A. Images of *Phyllosticta* species growing on a selection of different carbon sources. **B.** Clustering of *Phyllosticta* species based on their growth on different carbon sources. All species were grown on 35 different carbon sources, colony diameters were measured, and images takes on all sources when the biggest colony of a species reached the edge of its plate. All species grew fastest on wheat bran.

### 3.7 Genomes of *Dothideomycetes* can be clearly distinguished based on CAZyme content, but this does not correlate well with lifestyles described in literature

The genetic basis for the ability to utilize different carbon sources is often caused by differences in CAZymes repertoires (van den Brink and de Vries 2011; Lombard *et al*. 2014). Interestingly, the abundance and diversity of CAZymes encoded in a genome is also related to lifestyle and consequently enables to predict the tropic classification of a species (Lo Presti *et al*. 2015; Hane *et al*. 2020). To determine the trophic classification for *P. citrichinaensis*, we used CATAstrophy to annotate CAZyme genes for all 116 predicted proteomes and to perform a principal component analysis (PCA) to distinguish species with different trophic classes based on their CAZyme repertoire. CATAstrophy clearly separated species of different trophic classes based on the first principal component (PC1) (Fig. 5A and Suppl. Table 5). The second principal component (PC2) mainly separated species based on phylogeny; as previously observed for fungi and oomycetes (Hane *et al*. 2020), oomycetes clustered together separated along the PC2 axis from fungi. The different trophic classes differ considerably in the numbers of genes per CAZyme family (Fig. 5B). For example, GH families differ clearly between trophy class, and GH are also the CAZyme families with the highest number of genes per species. In contrast, almost no difference can be observed in the PL family, which generally has the smallest number of genes per species. The CATAstrophy gene predictions were used to identify CAZyme-containing OGs, for which we then obtained the number of genes present in each species to generate a clustered heatmap (Fig. 5C). We observed three distinct clusters that differ in their CAZyme repertoires, which typically correlate well with the CATAstrophy trophy predictions (Fig. 5C and Suppl. Table 5). The CATAstrophy trophy predictions also correlate well with the numbers of secreted proteins and effectors, although we did not observe such a strict separation into three clusters as was observed for CAZyme genes (Fig. 5C, Fig. 3).

**Figure 5.**
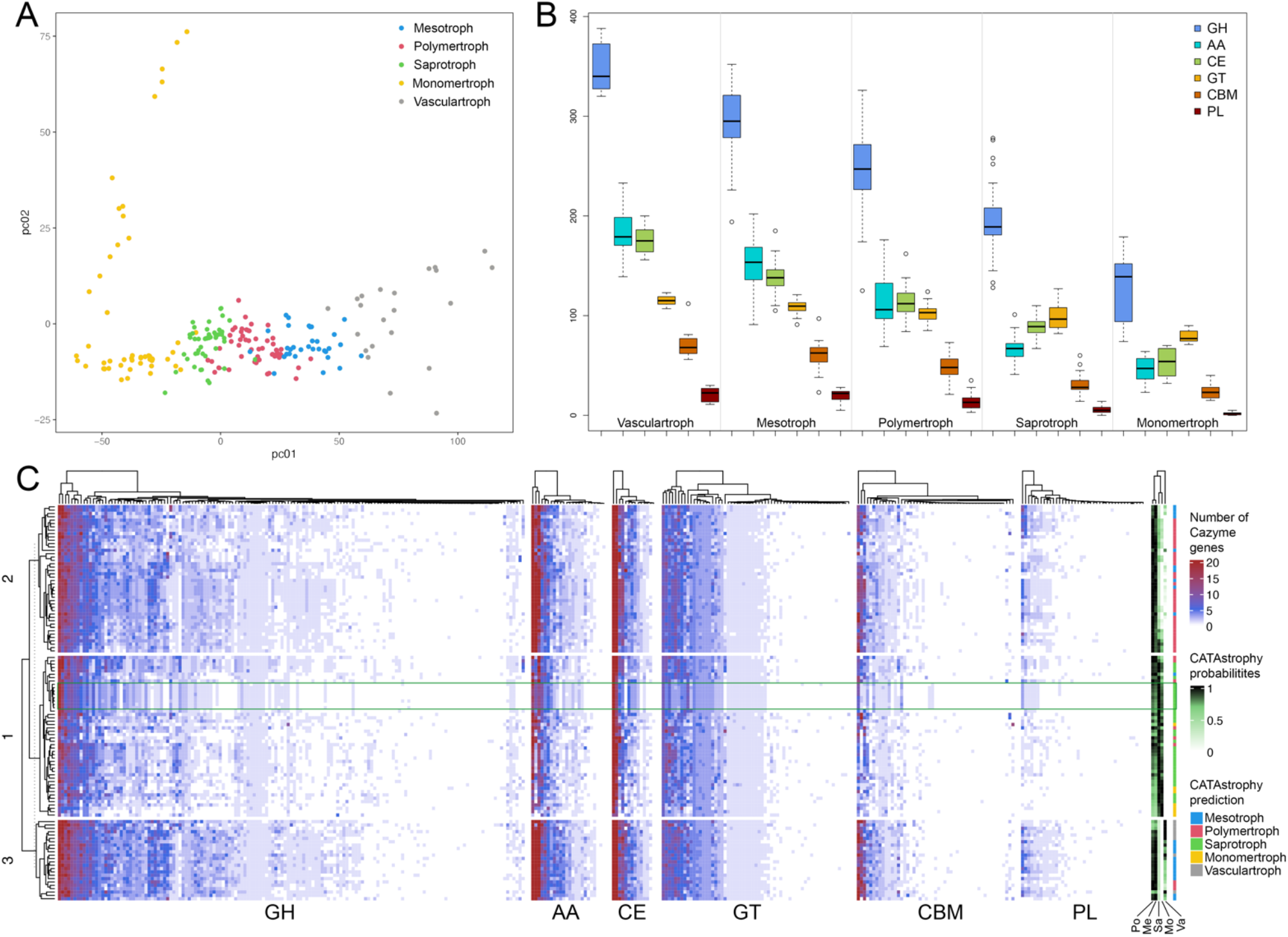
Separation of species into different trophy classes based on presence of CAZyme genes in their genomes. **A.** PCA plot of PC1 vs PC2. PC1 separates species of different trophy classes, PC2 separates species on phylogeny. **B.** Number of CAZyme genes in each CAZy family per CATAstrophy class. **C.** Clustered heatmap of the number of CAZyme-gene-containing OGs per species. GH = Glycosyl hydrolase, AA = Auxiliary activity, CE = Carbohydrate esterase, GT = Glycosyl transferase, CBM = Carbohydrate binding module, PL = Pectin lyase. Po = Polymertroph, Me = Mesotroph, Sa = Saprotroph, Mo = Monomertroph, Va = Vasculartroph.

All *Phyllosticta* species were predicted to be saprotrophs by CATAstrophy (Fig. 5C) and clustered according to their phylogenetic relationship, suggesting that they are generally very similar in terms of CAZymes (Suppl. Table 5). Consequently, *P. citrichinaensis* clustered most closely to *P. citribraziliensis*, an endophyte, as this is its closest relative (Fig. 1A). We nevertheless found six CAZyme families that show consistent differences in gene number between pathogens and endophytes: AA1_3, AA3, CBM18, CBM67, GH3, and PL22 (Table 1). In all cases except AA3, *P. citrichinaensis* follows the endophytic pattern. Together, these results indicate that in terms of the presence of CAZyme genes, the *P. citrichinaensis* genomes compare best to those of endophytes.

**Table 1.** CAZyme families with gene abundance differences between endophytes and pathogens. Columns with a darker fill color indicate the family which is more abundant in pathogens, while the lighter color indicates those which are more abundant in endophytes.

| Species name | Lifestyle | CBM67 | AA1_3 | CBM18 | GH3 | PL22 | AA3 |
| --- | --- | --- | --- | --- | --- | --- | --- |
| *Phyllosticta capitalensis* | E | 0 | 10 | 10 | 15 | 1 | 2 |
| *Phyllosticta citribraziliensis* | E | 0 | 10 | 9 | 15 | 1 | 2 |
| *Phyllosticta citrichinaensis* | ? | 0 | 10 | 9 | 16 | 1 | 1 |
| *Phyllosticta citrichinaensis* | ? | 0 | 10 | 9 | 15 | 1 | 1 |
| *Phyllosticta citriasiana* | P | 1 | 9 | 7 | 13 | 0 | 1 |
| *Phyllosticta citricarpa* (CPC 27913) | P | 1 | 9 | 6 | 14 | 0 | 1 |
| *Phyllosticta citricarpa* (CBS 127454) | P | 1 | 9 | 7 | 14 | 0 | 1 |
| *Phyllosticta paracitricarpa* | P | 1 | 9 | 7 | 14 | 0 | 1 |

## 4 Discussion

Lifestyle adaptations are thought to be driven by differences in gene content, and especially CAZymes are assumed to be crucial (ten Have *et al*. 2002; King *et al*. 2011; Hane *et al*. 2020). Here we aimed to elucidate the genomic differences between endophytes and pathogens within the *Phyllosticta* genus occurring on citrus, and aimed to determine the lifestyle of *P. citrichinaensis*. Based on the results we uncovered several differences between species with different lifestyles. For instance, endophytes more frequently contain higher numbers of genes in OGs, and these genes are more often annotated than in pathogenic species. In addition, pathogenic species share more un-annotated lifestyle-specific OGs as compared to endophytic species. Furthermore, we show that species cluster independently of phylogeny based on the CAZyme content of their genomes, and this clustering correlated well with trophy prediction by CATAstrophy. The ambiguous species *P. citrichinaensis* showed characteristics that matched with endophytes in some cases, with pathogens in other cases, and sometimes it did not match with either lifestyle, suggesting it may exhibit an intermediate lifestyle not accounted for in the current definitions.

We previously observed that only four CAZyme families showed a consistent difference between endophytic and pathogenic species (Buijs *et al*. 2021), one of which contained AA1_3/CBM18 (seven in pathogens vs eight in endophytes), which were mixed in one orthologue group, and another one contained family CBM18 only. In this study, we found in total six CAZyme families with a consistent difference between endophytes and pathogens. These included two separate OGs that contained the AA1_3 and CBM18 CAZyme families, which both consistently contained more genes in endophytes compared to pathogens. CATAstrophy predicts on average about 90 CAZyme genes extra compared to our previous results (Buijs *et al*. 2021). The biggest difference can be found in the CE (carbohydrate esterase) family, where between 83–89 genes are predicted instead of 15–17. Most of these are in the CE10 family (47–53). The CE10 family is no longer listed as carbohydrate active enzyme by the cazy.org database (used by JGI) because most of the members of this family act on non-carbohydrate substances (Lombard *et al*. 2014). If we manually remove these, CATAstrophy still predicts 20 extra genes in the CE family. In addition, there are some 15–20 extra genes predicted in the AA family, five to ten more in the GH family, and approximately five more in the GT family. Numbers in the PL family are practically identical. In the CBM family, CATAstrophy predicts about five to ten genes less. In the JGI annotation pipeline, many of the genes in the CBM family contained multiple domains and were therefore counted multiple times, which might have led to an overestimation. However, for most families it seems that CATAstrophy predicts more genes compared to JGI. As most of the CAZyme genes predicted by JGI are based on some experimental validation (cazy.org), it would be wise to experimentally validate the CAZyme genes that were predicted by CATAstrophy and not by JGI.

Genes related to secondary metabolite biosynthesis but not the number of biosynthetic gene clusters (BGCs) as predicted by antiSMASH differ between species. It is important to note here that BGC prediction by antiSMASH is based primarily on the presence of a gene to produce the metabolite ‘backbone’, such as a polyketide synthase (PKS) (Blin *et al*. 2019). Once such a gene is found, antiSMASH takes an area in the genome of up to 20 kb (depending on the type of backbone gene) on either side of the gene and checks for the presence of tailoring genes, which are then all automatically included in the BGC. This means that tailoring genes that are altered or inactive are still included in the predicted BGC, and that alterations in genes may not result in an altered BGC prediction. In contrast, although the cluster may look very similar, alterations in tailoring genes may lead to the production of a very different compound: a good example is the synthesis of the toxins dothistromin, aflatoxin, and sterigmatocystin, which are all synthesized in a very similar manner, with only the very last tailoring steps being different (Schwelm and Bradshaw, 2010; for a review see Hüttel and Müller, 2021). We therefore conclude that although antiSMASH did not detect differences in BGC numbers, alterations in biosynthetic genes may be responsible for the differences between species of different lifestyles in the genus *Phyllosticta*, and this will be an interesting subject for future studies.

CATAstrophy was able to clearly separate species based on the number of CAZyme genes present in the genomes and was able to separate species into different trophy predictions. Apart from closely related species that cluster together, a phylogenetic pattern cannot be observed in the clustering of the heatmap or in the trophy predictions, which suggests that there is a strong signal that links genome content to lifestyle. CATAstrophy also allows for trophy-overlap for species that are bordering two trophies, such as for *Sclerotinia sclerotiorum*, which received very high scores for both the polymertroph and the saprotroph class (Suppl. Fig. 5). This is consistent with literature, as *S. sclerotiorum* has been described to exhibit a necrotrophic phase that is followed by a saprotrophic phase (Hegedus and Rimmer 2005). *Sclerotinia sclerotiorum* is a much-researched model organism, and the descriptions in literature of its lifestyle are therefore well-developed. However, for many fungal species, this is not the case, and circumscriptions in literature are often limited or conflicting. Indeed, we see that the trophy predictions by CATAstrophy do not always correlate well with lifestyles described (or supposed) in literature: each trophy class includes species that are described as pathogens, endophytes, symbionts, or saprotrophs, or which have been described to exhibit multiple of these lifestyles. An underlying cause for this fact is that lifestyles that are described in literature may be inaccurate as the border between species of different lifestyles such as necrotrophs, hemibiotrophs, or biotrophs is not very strict, or need very specific conditions to manifest. For instance, *Phytophthora infestans* has been placed in all three classes (Oliver and Ipcho 2004). Similarly, *Botrytis cinerea* has been placed in different classes as the symptoms it causes differ widely in their severity, depending on the exact interaction with the host (Veloso and Van Kan 2018). Within the genus *Phyllosticta*, some obscurity with respect to lifestyle is present for instance for *P. capitalensis*, which is a widespread endophyte of many hosts including citrus (Wikee *et al*. 2013a), but may cause disease in other hosts such as guava (Arafat 2018). In addition, non-pathogens can evolve a pathogenic lifestyle, and vice-versa, as can be observed with pathogenicity on pea by *Neocosmospora solani*, which is dependent on the presence of only a few genes, or with pathogenicity of *Fusarium oxysporum* on cucurbit species, which is determined by the absence or presence of a mobile pathogenicity chromosome (Temporini and Vanetten 2002; Dong *et al*. 2015; van Dam *et al*. 2017; Möller and Stukenbrock 2017). The possibility for species to be categorized into multiple trophies in the CAZyme-based classification system therefore presents an advantage over the traditional classification system as it allows for a more correct, double classification of species that exhibit multiple lifestyles depending on the host and other environmental parameters.

We compared genomes of species with different lifestyles within the genus *Phyllosticta* specifically, and found several distinctions. We observed that endophytes more often have higher numbers of genes in specific OGs as compared to pathogens. This suggests that the ability to be an endophyte necessitates the presence of additional genes. The ancestral *Dothideomycete* was likely a saprotroph, however, the most common ancestor of the *Botryosphaeriales*, the order in which *Phyllosticta* resides, was probably a plant pathogen, as determined by ancestral state reconstruction (Abdollahzadeh *et al*. 2020; Haridas *et al*. 2020). The evolution of *Phyllosticta* endophytes from *Phyllosticta* pathogens through a gain of genes and thereby gain of abilities is therefore a plausible scenario. With respect to lifestyle definition in *Phyllosticta*, *P. citrichinaensis* is the most ambiguous citrus-related species within the genus. By comparing the genome of this species with those of other species with different lifestyles within the genus *Phyllosticta*, we aimed to elucidate the lifestyle of this species. In several aspects, we found *P. citrichinaensis* to be most similar to endophytes. For instance, *P. citrichinaensis* shares more OGs specific to species of one lifestyle with endophytic species (50) than it does with pathogenic species (30), and none of its biosynthetic gene clusters was predicted to produce a squalestatin, which was also the case for all of the endophytes, but not for the pathogens. In addition, CAZyme families that had more genes in endophytes than in pathogens, often also had more genes in *P. citrichinaensis*. Furthermore, growth of *P. citrichinaensis* was not inhibited by the presence of sugar beet pulp, similarly to endophytic species. In contrast, its broader carbon utilization capabilities were more comparable to those of pathogenic species. Another aspect in which *P. citrichinaensis* was comparable to pathogens, was the presence of more putative peroxiredoxin genes in pathogens and *P. citrichinaensis* as compared to endophytes. On other aspects, *P. citrichinaensis* did not compare well with species of either lifestyle, such as the number of OGs that had more genes in species of either lifestyle: in almost a third of the cases, *P. citrichinaensis* did not match the gene numbers of either lifestyle. The number of effector genes that *P. citrichinaensis* shared with species of either lifestyle also suggests an intermediate pattern. The lifestyle of *P. citrichinaensis* cannot be univocally determined without performing pathogenicity assays, but as these are currently not available for this species, the data presented here give a good estimation that shows that *P. citrichinaensis* is an intermediate taxon, not perfectly fitting into any of the currently defined lifestyle definitions.

Research performed in recent years has shown with increasing confidence that borders between lifestyles simply are not very strict and in fact are subject to constant change. Examples such as *B. cinerea* and *Phytophthora infestans*, which both have been placed in multiple lifestyle classes depending on the host and other environmental parameters, demonstrate that our current classification systems are not always adequate to separate species into different lifestyles (Oliver and Ipcho 2004; Veloso and Van Kan 2018). In addition, the ability of species to be pathogenic to a specific host may change with the gain of only a few genes (Temporini and Vanetten 2002; van Dam *et al*. 2017). Classifying plant-associated microbes into different lifestyles is an important area of research as it allows for the identification of genomic parameters that are required for pathogenicity, and therefore aids in the search of a remedy against such pathogens. However, a classification system is only valuable if it allows for the accurate separation of species; an incorrectly classified organism may lead to incorrect conclusions and could cause much confusion. A classification such as the one proposed by Hane *et al*., which is based on the number of CAZyme genes in a species’ genome, is a significant improvement since it allows for overlap between lifestyles. Further development of such classifications for instance by the addition of other genomic parameters such as the presence of effectors could lead to the development of a more accurate and useful classification system in the future.

## Data availability statement

The whole genome sequencing data including annotations for the newly sequenced *Phyllosticta citrichinaensis* genome are publicly available at the JGI genome portal: https://mycocosm.jgi.doe.gov/Pcit129764. The authors affirm that all other data necessary for confirming the conclusions of the article are present within the article, figures, and (supplementary) tables.

## Acknowledgements

This work was funded by the Dutch Applied Science division (TTW) of NWO and the Technology Program of the Ministry of Infrastructure and Water Management under project 15807 of the Research Programme I&W Biotechnology and Safety. The work conducted by the U.S. Department of Energy Joint Genome Institute, a DOE Office of Science User Facility, is supported by the Office of Science of the U.S. Department of Energy under Contract No. DE-AC02-05CH11231.

## Supplementary Figures

**Supplementary Fig. S1.**
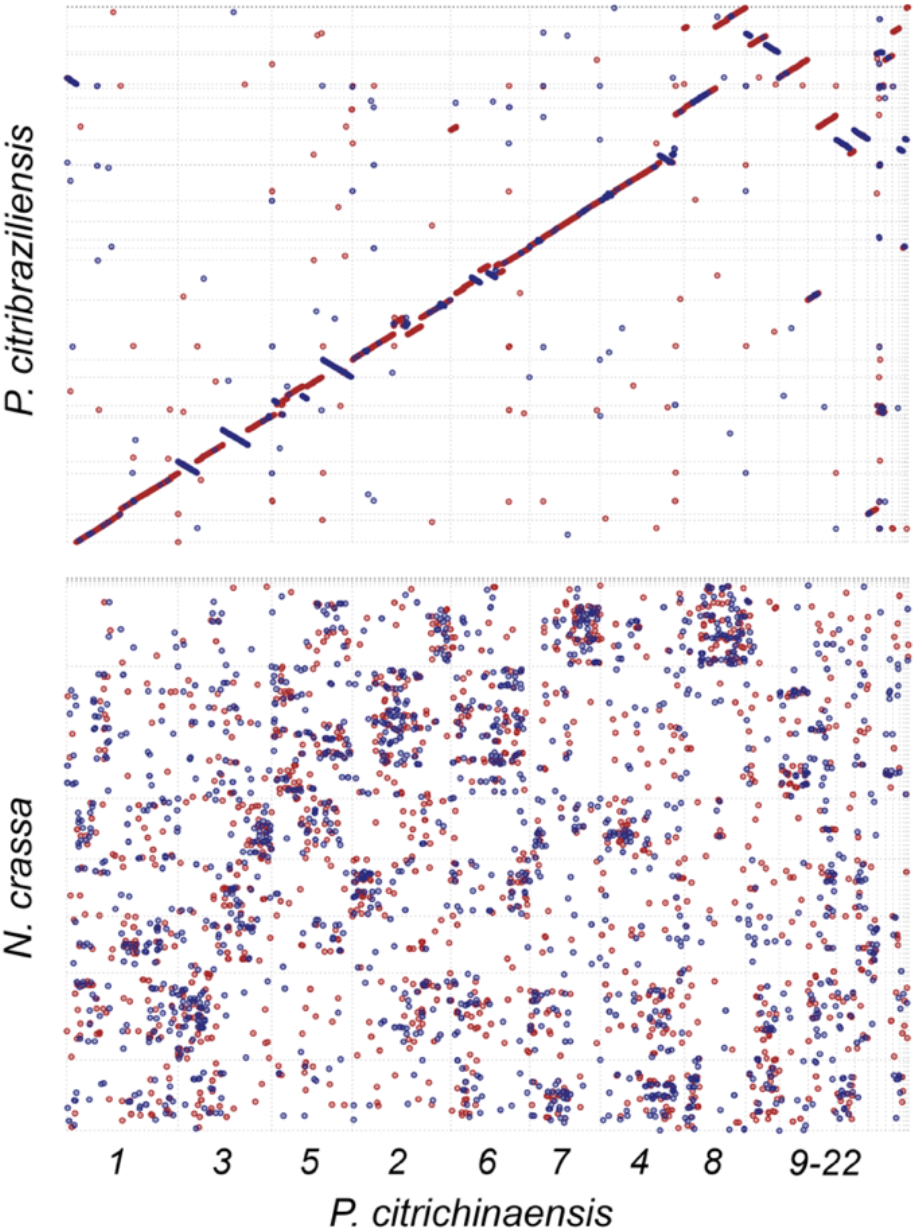
Whole-genome alignments between *P. citrichinaensis* and *P. citribraziliensis* (top panel) and *Neurospora crassa* (lower panel) were generated using PROmer. The red line/dot = homologous area and the blue line/dot = reversed homologous area.

## Supplementary Tables (submitted as separate excel files)

**Supplementary Table 1. Genome data**, including QUAST and BUSCO results, of all species used in this study.

**Supplementary Table 2. Orthofinder results. A**. Orthofinder statistical data. **B**. List of ortholog groups (OGs) that are unique in species of one lifestyle. **C**. List of OGs that have more genes in species of one lifestyle.

**Supplementary Table 3. Annotation data of OGs. A.** Annotation data of OGs present only in endophytes. **B**. Annotation data of OGs present in endophytes and *P. citrichinaensis* (PCC). **C**. Annotation data of OGs present only in pathogens. **D**. Annotation data of OGs present in pathogens and *P. citrichinaensis* (PCC). **E**. Annotation data of OGs that consistently contain more genes in *Phyllosticta* endophytes as compared to *Phyllosticta* pathogens. **F**. Annotation data of OGs that consistently contain more genes in *Phyllosticta* pathogens as compared to *Phyllosticta* endophytes. **G**. Annotation data of OGs in which the gene number in *P. citrichinaensis* (PCC) matches the number of endophytes. **H**. Annotation data of OGs in which the gene number in *P. citrichinaensis* (PCC) matches the number of pathogens. **I**. The number of OGs in each lifestyle-related group that is classified to each KOG class. **J**. A list of annotation files from the JGI database that were included in these analyses.

**Supplementary Table 4. Data of OGs that contain putative secreted and effector proteins. A.** Secreted OGs in listed in order corresponding to Figure 3A. **B**. Effector OGs listed in order corresponding to Figure 3B. **C**. Number of secreted genes as estimated by: number of genes in secreted OG vs signalP prediction. **D**. Number of effector genes as estimated by number of genes in effector OG vs effectorP prediction. **E**. Effector OGs that are unique to species of *Phyllosticta*. **F**. Effector OGs that are present in all *Phyllosticta* spp. **G**. Functions of secreted OGs that are unique to *Phyllosticta*.

**Supplementary Table 5. CATAstrophy results.** Including all trophy predictions and all CAZyme gene counts.

